# COVID-19-associated pulmonary aspergillosis in immunocompetent patients: A virtual patient cohort study

**DOI:** 10.1101/2022.07.18.500514

**Authors:** Henrique AL Ribeiro, Yogesh Scindia, Borna Mehrad, Reinhard Laubenbacher

## Abstract

**Purpose:** The opportunistic fungus *Aspergillus fumigatus* infects the lungs of immunocompromised hosts, including patients undergoing chemotherapy or organ transplantation. More recently however, immunocompetent patients with severe SARS-CoV2 have been reported to be affected by COVID-19 Associated Pulmonary Aspergillosis (CAPA), in the absence of the conventional risk factors for invasive aspergillosis. This paper explores the hypothesis that contributing causes are the destruction of the lung epithelium permitting colonization by opportunistic pathogens. At the same time, the exhaustion of the immune system, characterized by cytokine storms, apoptosis, and depletion of leukocytes may hinder the response to *A. fumigatus* infection. The combination of these factors may explain the onset of invasive aspergillosis in immunocompetent patients.

**Methods:** We used a previously published computational model of the innate immune response to infection with *Aspergillus fumigatus*. Variation of model parameters was used to create a virtual patient population. A simulation study of this virtual patient population to test potential causes for co-infection in immunocompetent patients.

**Results:** The two most important factors determining the likelihood of CAPA were the inherent virulence of the fungus and the effectiveness of the neutrophil population, as measured by granule half-life and ability to kill fungal cells. Varying these parameters across the virtual patient population generated a realistic distribution of CAPA phenotypes observed in the literature.

**Conclusions:** Computational models are an effective tool for hypothesis generation. Varying model parameters can be used to create a virtual patient population for identifying candidate mechanisms for phenomena observed in actual patient populations.

## 1 Introduction

*Aspergillus fumigatus* is an opportunistic fungus that can infect the lungs of immunocompromised hosts, including, among others, patients undergoing chemotherapy or receiving an organ transplant, and patients affected by chronic granulomatous disease [1, 2]. Neutrophils are essential to fighting the pathogen, and impaired neutrophil responses are a common predisposing factor to the infection. More recently, patients with severe SARS-CoV2 infection have reportedly been affected by COVID-19 Associated Pulmonary aspergillosis (CAPA). However, Lai, CC, and Yu, W 2021 [3] found that the conventional risk factors for aspergillosis were not present in CAPA patients.

Mitaka, H *et al*. 2021 [4] found that around 10% of COVID-19 patients in ICUs develop invasive aspergillosis, and the mortality rate for these patients is 54% compared with 24% of those without CAPA [5]. According to Lai, CC, and Yu, W 2021 [3], patients with CAPA are predominantly male, with an average age of 73±13 years, and 88% had diabetes, high blood pressure, kidney disease, chronic obstructive pulmonary disease (COPD), or heart disease. The fact that CAPA is associated with these comorbidities but not with the conventional risk factors for invasive aspergillosis requires the generation of new hypotheses to explain the susceptibility of these hosts to the infection.

Computational models have been used extensively in immunology to create and test hypotheses. In particular, agent-based models have proven effective for studying respiratory diseases. They are intuitive, rule-based models that make it easy to represent heterogeneous spatial environments and individual entities, such as immune cells or pathogens, and their generation does not require extensive modeling expertise. We published an agent-based model of the innate immune response to *Aspergillus fumigatus* within an alveolar duct that established that one of the critical parameters determining infection outcome is the distance within which macrophages can detect fungal spores [6].

In [7], we published an agent-based model of the immune response to *Aspergillus fumigatus.* The model focuses on the role of iron sequestration by the host as part of the innate immune response. It was parametrized entirely with information from the literature rather than data fitting, and was extensively validated with both data from the literature and our own time course data from experiments using a mouse model of the infection. We also showed that variation of parametrizations from this reference model could account for the variability across experiments. A first result we obtained using the model demonstrates that fungal strains engage different parts of the innate immune response depending on their level of virulence. This computational model is the main tool for the results reported here.

We hypothesize that in patients with severe COVID-19, the destruction of the lung epithelium facilitates colonization by opportunistic pathogens. At the same time, the exhaustion of the immune system characterized by cytokine storms, apoptosis, and depletion of leukocytes hinders its ability to kill the fungus, explaining the susceptibility of otherwise immunocompetent patients to aspergillosis. We use the computational model to test this hypothesis and to ask what set of parameters and conditions explain CAPA. To test our hypothesis, we created a virtual patient population by sampling the model parameter space, characterizing patients with dual infection and analyzing disease outcomes, with each choice of parameters representing one virtual patient.

## 2 Material and Methods

### 2.1 Modeling

The simulation study in this paper uses a computational model of the coinfection that is a modification of our previously published model [7] capturing key features of the innate immune response to a respiratory *Aspergillus fumigatus* infection in immunocompetent hosts, including the role of iron regulation, an important virulence factor. The model spans the intracellular, tissue-level, and organism scales. At the tissue level, an agent-based model simulates the immune response in a spatially homogeneous representation of a volume of lung tissue. The domain simulated (6.4 × 10^-2^ *μ*L) is enough to represent a piece of lung infected with a few conidia. Its components are depicted in Figure 1. The original model of immune response to *Aspergillus fumigatus* had 76 parameters; here, we use a reduced version of that model with 27 parameters (Table A.2) by eliminating the iron regulation component and aggregating some of the cytokines represented explicitly. We describe the model in detail below.

**Fig. 1.**
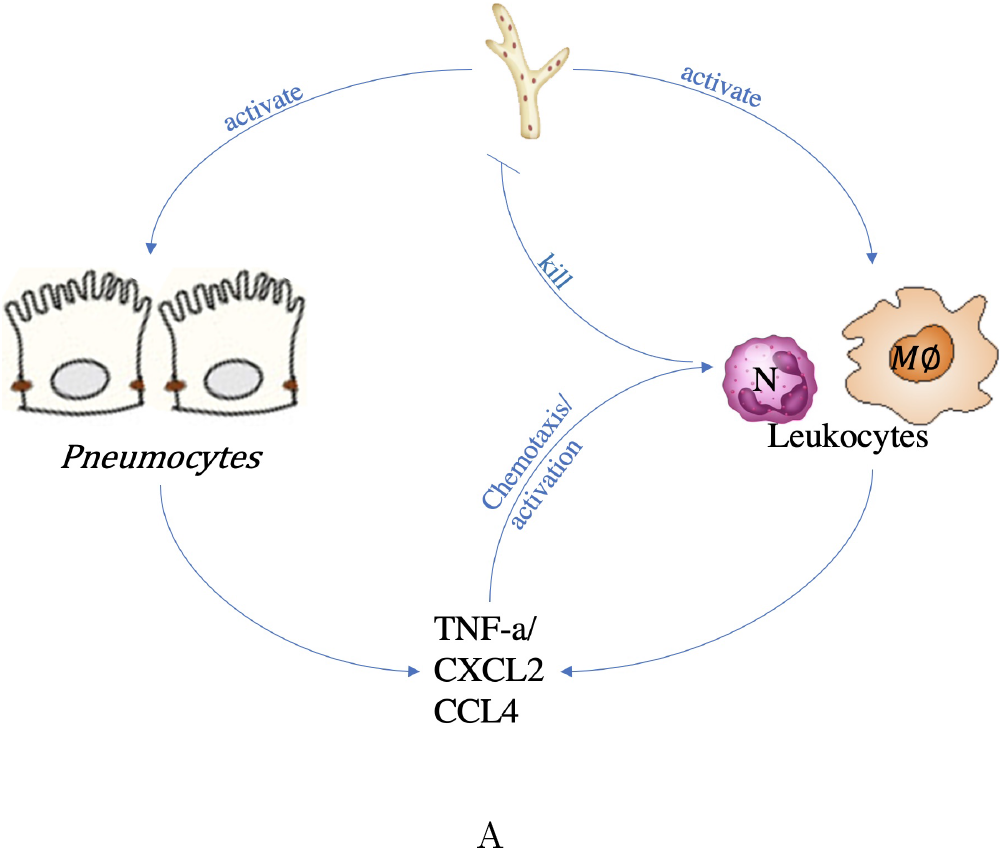
Model description. *Aspergillus fumigatus* hyphae activate host cells (pneumo-ytes, monocytes, and neutrophils). Pneumocytes and monocytes secrete cytokines (TNF and IL-10) and chemokines (CXCL2 and CCL4). The chemokines chemoattract other leukocytes (neutrophils and monocytes) to the infection site, and the cytokines (TNF), together with the fungus itself, activate the leukocytes. Leukocytes kill *A. fumigatus*. This simplified figure does not show inactivation by IL-10/TGF-*β*/apoptotic cells and the secretion of TGF-*β* by inactivated cells.

The 3-dimensional space is homogeneous with periodic boundary conditions; that is, a 3-dimensional torus. If a molecule or agent leaves the simulation space by crossing one boundary, it re-enters from another, similar to previous modeling in this context (e.g., [8, 9]). The rationale for periodic boundary conditions is that the simulation covers a small volume amid a large infected volume. Therefore, the concentration of molecules across all boundaries should be similar.

As in our published model, we have three types of host cells: type II epithelial cells, macrophages, and neutrophils. These cells are equipped with a simple intracellular model (Figure 2) that determines their state at any given time. They receive signals in the form of cytokines or contact with *A. fumigatus* and move to one of three final states: Active (macrophages, epithelial cells, and neutrophils), TNF primed (macrophages and epithelial cells), or Inactive (macrophages only). Each state allows the cell to engage in certain activities, such as secreting specific cytokines. Table A.1 shows the interactions between agents and molecules and the outcomes of these interactions. It is worth noting that only active macrophages can kill hyphae, while neutrophils in any state can kill. That was an assumption in our previous model, based on evidence that macrophages need pre-activation to kill hyphae [10, 11].

**Fig. 2.**
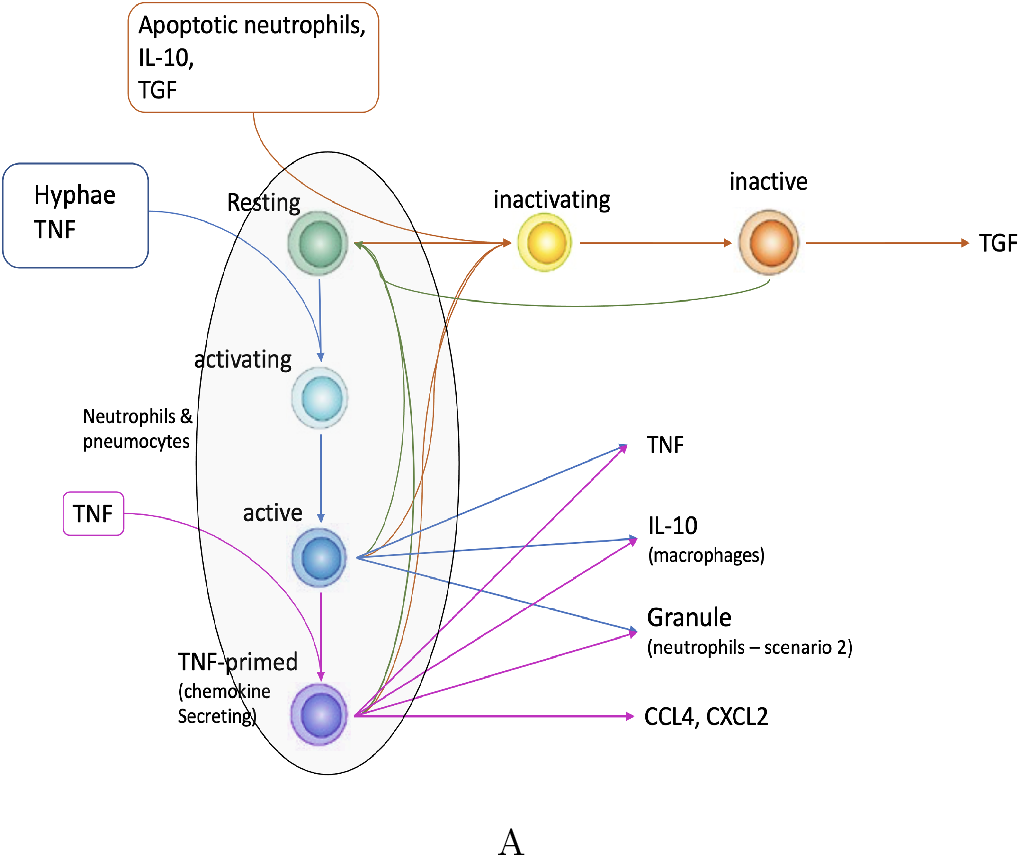
Diagram showing host cell state changes. In response to TNF or hyphae, resting host cells move to the activating state, and after 60 iterations activating cells become active. Active cells extra primed with TNF become chemokine secreting. Macrophages primed with TGF-*β*, IL-10, or apoptotic cells become “inactivating” and then inactive (TGF-*β* secreting). After 180 iterations, inactive, active, or TNF-primed cells return to the resting state without extra signaling. The times to move from one state to another are approximations using the times for gene activation and inactivation measured in *in-vitro* experiments [7].

Figure 2 and Table A.1 summarize most features of the model dynamics. As mentioned, in this model we do not include iron metabolism explicitly, thereby significantly reducing the number of parameters. We also assumed that there were no significant changes in cell counts over time, due to the fact that we only simulate a 24 hour period, based on the observation in [12] that the number of monocytes and neutrophils is approximately constant from day two to three, which makes our assumption reasonable. We used two minute time steps in order to correctly capture the dynamics of diffusion, since a longer time step would lead to near equilibrium in the diffused molecules.

Besides cells, the model also contains five molecules: IL10, TNF, TGF-*β*;, CCL4, and CXCL2. These molecules diffuse through the space according to a partial differential equation [13] and interact with cells with probability given by Equation 1:

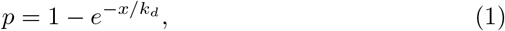

where *p* is the probability that the receptor will be activated, *x* is the cytokine concentration, and *k_d_* is its dissociation constant. This equation is also used for the reaction between granules and hyphae. In this case, *x* is the granule concentration in arbitrary units, *k_d_* is *k_d_*_GRANULE, and *p* is the probability that the hyphal septae will die. In the case of hyphae killing by granules this equation is a phenomenological approximation. Moreover, a given molecule’s concentration decays with a half-life of one hour and a continuous exchange with the serum (Equation 2):

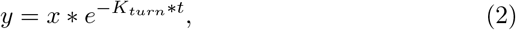

where *K_turn_* is the turnover rate, and *t* is the time-step size (2 minutes).

The two mechanisms not covered so far are leukocyte movement and *A. fumigatus* growth. Like in our published model, in the absence of chemokines, cells move randomly, while in their presence, they tend to move to the voxels with higher chemokine concentrations. The rate of movement is constant, and the cells will, on average, traverse a fixed number of voxels per time step. In the presence of chemokines, each voxel receives a weight according to Equation 3, where CCL4 guides macrophages and CXCL2 guides neutrophils [7]:

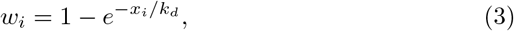

where *W_i_* is the weight of the *i*-th neighbor voxel, *x_i_* is the chemokine concentration in the *i*-th voxel, and *k_d_* is the dissociation constant.

In nature, hyphal growth is of course a continuous process composed of elongation and branching (chiefly sub-apical branching). In the model we represent hyphal growth and branching as a discrete approximation. Every 40 *μm* of septae was considered a unit (we called it a cell in the simulator). A tip cell can produce another tip cell (elongation), while a sub-tip cell can form a 45° branch (subapical branch) [14, 15] with a 25% probability. Other cells cannot originate new cells unless their neighbors are killed and they become tip or sub-tip cells again. In this paper, we did not change the branching probability, and to vary the growth rate, we changed the time it takes for a tip cell to generate a new unit.

### 2.2 Modeling Coinfection

We model the COVID-19 *A. fumigatus* coinfection implicitly by changing specific parameters as well as the number of leukocytes in the lung space (Table 1). In the model, neutrophils kill hyphae by direct contact. However, to accommodate alternative mechanisms of neutrophil anti-microbial function, we introduced a second scenario in which leukocytes secrete granules that diffuse and kill the fungus. In our published model, we considered neutrophils killing *A. fumigatus* via ROS secretion only. The ROS concentration to kill hyphae is high enough only in the synapsis between the neutrophil and hyphae. However, other granule molecules may also be able to kill hyphae, hence the second scenario to account for uncertainty in the mechanism.

**Table 1.** Model parameter ranges. The table shows the ranges in which we sample parameters (maximum and minimum value). Column 1: parameter name; Column 2: description; Column 3: parameter range; Column 4: parameter reference value; Column 5: rationale for the value in Column 4. The values taken from [7] are the default values in our previous work.

| Parameter | Description | Parameter range | Reference Value | Ref |
| --- | --- | --- | --- | --- |
| GROWTH_RATE | intrinsic growth rate | 2.75-80 $\mu\text{m}/h$ | 40 $\mu\text{m}/h$ | [7] |
| PR_N_KILL | probability of neutrophils to kill hyphae | 0.23-23% | 23% | [7] |
| PR_MA_KILL | probability of monocytes to kill hyphae | 0.099-9.9% | 9.9% | [7] |
| NUM_MA | number of monocytes | 360-960 cells | 960 cells | [16] |
| NUM_N | number of neutrophils | 360-960 cells | 360 | [16] |

The results for the second scenario are presented in the supplementary material. For the second scenario, we introduced the parameters (*k_d_*_GRANULE and GRANULE_HALF_LIFE - Table A3). Upon interaction with *A. fumigatus*, neutrophils are activated and then secrete one arbitrary unit of granule (an abstraction for the molecules contained in neutrophil granules). This granule kills hyphae with a probability given by Equation 1. The granule contents, like all molecules, decay with a given half-life (GRANULE_HALF_LIFE). The rationale for using arbitrary units is simple. All we need is a pair of values, GRANULE_QTTY and *k_d_*_GRANULE, that fits the data in [7]. This pair of parameters is not identifiable, so we fixed GRANULE_QTTY to one arbitrary unity and fit the *k_d_* to reproduce the simulation data in our previous work.

In [7], iron sequestration acted to inhibit *Aspergillus fumigatus* growth. Here we do not consider the role of iron explicitly. Instead, we tested a wide range of hyphal growth rates (Table 1) to represent more or less permissive lung environments, partly as the result of variable availability of nutrients such as iron across the space (see below). Another contributing factor is the damage done to the epithelium by the ongoing viral infection.

In this study, we focus on five parameters, with values listed in Table 1. The first parameter is the intrinsic growth rate of the fungus. The second and third parameters are proxies for the strength of the immune response, and the fourth and fifth parameters reflect an increased number of immune cells as part of the response to the viral infection. The intrinsic growth rate of the hyphae encapsulates the permissivity of the environment, such as the nutrient availability. However, the collective effects of the intrinsic growth rate, together with the counteracting effects of the immune response, we call the observed growth rate.

We performed Latin-Hypercube Sampling (LHS) on the parameters in Table 1. We varied the number of leukocytes, the monocyte killing probability (the probability with which monocytes kill hyphae upon direct contact), the intrinsic growth rate, and the neutrophil killing probability (the probability with which neutrophils kill hyphae upon direct contact). The intrinsic growth rate varies between 2.75 *μm/h* to 80 *μm/h.* The rationale for these values is that the host limits access of the fungus to nutrients, thereby decreasing its growth rate. Therefore, at the lowest restriction, the fungus will grow to its full potential. The literature measure *A. fumigatus* growth *in vitro* of 40 *μm/h* (see our previous work [7]) and 60 *μm/h* [12]. Because there is uncertainty in these numbers and in-vitro growth may be conservative, we set the maximum to 80 *μm/h*.

The probability of neutrophils and macrophages killing hyphae had a similar rationale. We had the default from our previous work of 23% and 9.9%, respectively. Because we are considering the negative effect of SARS-CoV-2 in these cells (See Results and Discussion), these numbers can only decrease. Meanwhile, the number of leukocytes reflects the uncertainty and natural variability in these cells. We use the ranges given by [12] (5-15 million) as an educated guess. Note that at the peak, on day one, we have 15 million neutrophils, and on day three, 15 million monocytes. Therefore, these ranges are reasonable for both cell types.

Using LHS sampling we generated 24,000 parameter sets, representing 24,000 virtual hosts of the dual infection. (We also created an additional 36,000 virtual hosts using the alternative neutrophil mechanism; see supplementary material). We measured the observed growth rate as the slope of the log of the *A. fumigatus* curve across the 24h of simulation (Equation 4). Note that we shifted the *A. fumigatus* curve by 1 to avoid negative infinity when the number of *A. fumigatus* cells tends to zero. The observed growth rate is a value given by Equation 4, while the intrinsic growth rate is a model parameter (Table 1). In Equation 4 we are counting the number of hyphal septae:

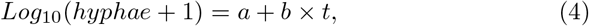

where *b* is the observed growth rate, and *t* is time in hours.

### 2.3 Partial Rank Correlation Coefficient

A partial rank correlation coefficient is a common way to assess local model sensitivity to parameters. In this paper, we performed a sensitivity analysis (SA) of the five (Scenario 1) and six (Scenario 2) parameters against *A. fumigatus* observed growth rate. We used the method “pcc” from the R package “verification” [17]. To do the SA we used our virtual patient population.

Moreover, we also measured how the parameters’ partial correlation changed as the intrinsic growth rate changed. That is, we divided the virtual patient population into ten bins of similar intrinsic growth rates (GROWTH_RATE). Then we computed the partial rank correlation coefficient of the other four parameters (PR_N_KILL, PR_MA_KILL, NUMMA, and NUMN).

### 2.4 Code

The simulator was written in Java (JavaSE-1.8) and needs only the JRE System Libraries. A typical execution takes about 4-5 seconds to simulate 24 hours in an OSX 2.6 GHz 6-Core Intel Core i7. Code is available at: https://github.com/deassisinfo/CAPA/

## 3 Results

Our goal is to test the hypothesis that a respiratory viral infection such as severe COVID-19 can make the lung environment more permissive to *A. fumigatus* growth and, at the same time, exhaust the immune system, allowing the fungal infection to progress in an otherwise immunocompetent host. In the simulator, these conditions translate into a higher intrinsic growth rate and lower ability of leukocytes to kill hyphae. Concomitantly, when the *A. fumigatus* infection starts, the number of leukocytes in the lung is high because the host is already fighting another infection. These conditions are encoded by the model parameters in Table 1. Each parameter sample can be thought of as a virtual patient for whom we simulate infection outcome. We started each simulation with 640 type-II pneumocytes, 20 germinated conidia, and 360-960 monocytes and neutrophils. Each simulation represented 24 hours (720 iterations), and we used a time-step of 2 minutes. The results are shown in Figure 3.

**Fig. 3.**
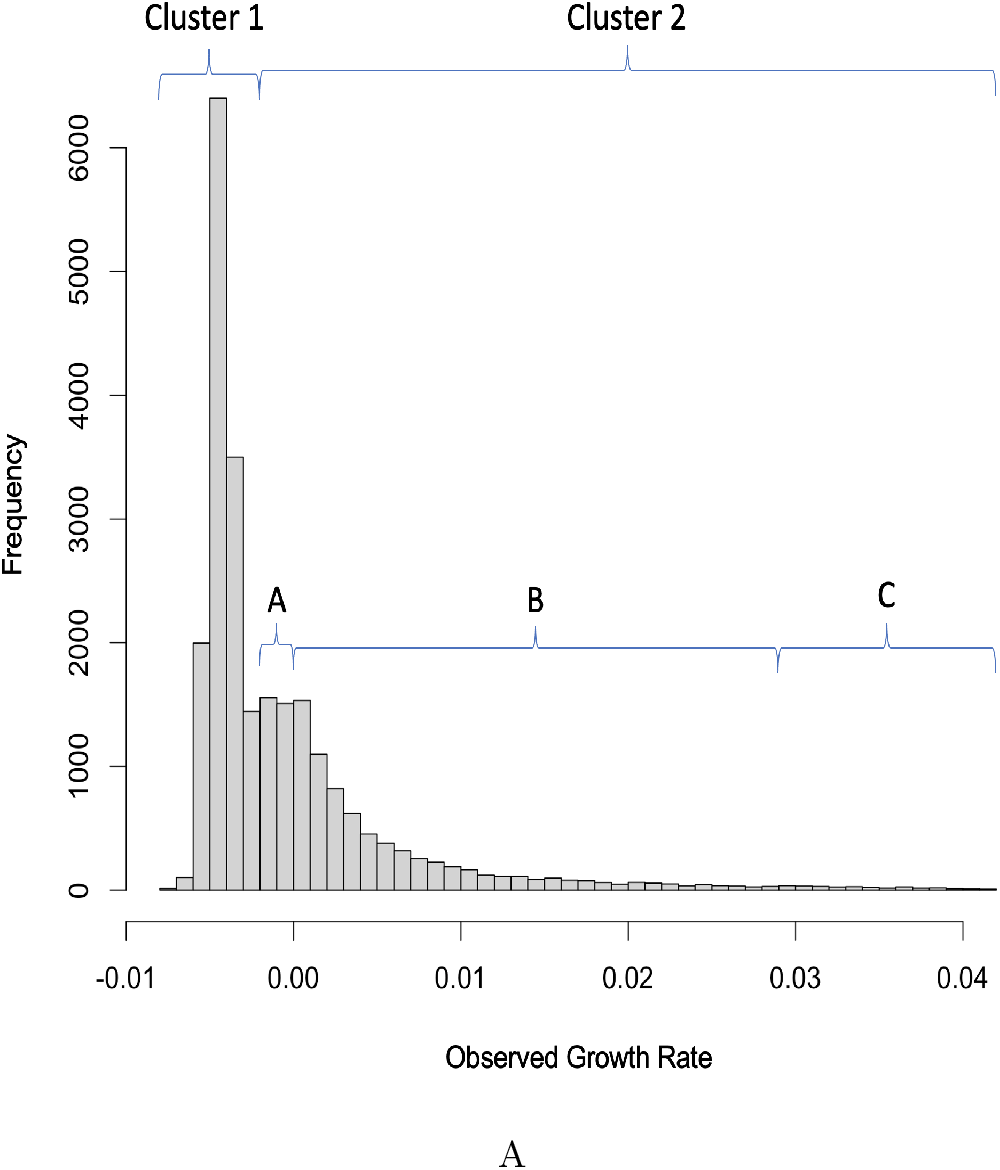
Distribution of observed growth rate. The observed growth rate of *A. fumigatus* was measured during the 24 hours of simulation in the virtual patient population. The graph shows a bimodal distribution of the growth rate. We divided the virtual patients into two clusters according to their observed growth rate: the prominent peak on the negative side (Cluster 1) and the secondary peak (Cluster 2). We then subdivided Cluster 2 into three subclusters: 2A is the negative part, 2B is the positive part excluding outliers, and 2C is the long tail of outliers.

In order to correctly interpret Figure 3, one needs to keep in mind the distinction between intrinsic growth rate and observed growth rate. The first is a parameter (Table 1) that controls the time it takes for new *A. fumigatus* cells to be generated. The second results from the number of cells generated (intrinsic growth rate) minus the number of cells killed by leukocytes. The intrinsic growth rate is always positive, while the observed growth rate may be negative if the infection is declining, as observed in the vast majority of cases in Figure 3.

Observe that the distinction between intrinsic and observed growth rates is exclusively a model property. The intrinsic growth rate is a model parameter, whereas the observed rate is a model output. However, from the point of view of nature or a more complex model, what we call intrinsic may still be considered observed. For example, a simple model may have a growth rate with a value of 3 *μm/h*. Therefore, for this model, 3 *μm/h* is its intrinsic growth rate. However, a more complex model may have two parameters:’ growth rate and iron acquisition, none of which are 3 um/h. But they interact to produce an observed growth rate of 3 *μm/h*.

The data in Figure 3 show a prominent peak in negative observed growth rate, indicating resolution of the infection, and a second smaller peak with a long tail of positive observed growth rate. We can therefore cluster patients into two groups, with Cluster 1 containing only patients that clear the infection. Cluster 2 can be further subdivided into three subclusters representing progressively worsening fungal infection. Cluster 2A is the portion of Cluster 2 with negative observed growth rate, representing patients whose infection clears, while Cluster 2B is the portion of Cluster 2 with positive observed growth rate (excluding the outliers), and Cluster 2C is the long tail of outliers (Figure 3), both together representing patients that develop CAPA. We found a similar distribution in Scenario 2 (Figure A2). This bimodal distribution is consistent with what is known for this disease: patients with an intact neutrophil response resolve the fungal infection, whereas the cluster with positive observed growth rate corresponds to the observation that COVID-19 patients in the ICU can develop CAPA [4]. Figure A1 shows that the percentage of patients with CAPA in our simulation (clusters 2B and 2C) is similar to the one measured in epidemiological studies.

Clusters 2B and 2C in Figure 3 and Table A4 suggest that a combination of higher intrinsic fungal growth rate and lower fungal killing by the immune response can explain the onset of CAPA in otherwise immunocompetent hosts. Figure A4 confirms that patients with CAPA have decreased ROS production by neutrophils, one of the key tools used by leukocytes to kill hyphae. This reduction is qualitatively similar to the reduction in neutrophil-killing ability between CAPA and non-CAPA patients (Figure A4B). Meanwhile, Figure A5 shows that hemorrhage and the availability of heme is a factor that favors *A. fumigatus* growth, similar to the increase in the observed growth rate observed in our CAPA virtual patients (Figure A5C). Interestingly in Scenario 2, the intrinsic growth rate seems to play a secondary role (Table 2). To further explore this hypothesis, we constructed classification trees to see which combinations of parameters are necessary for CAPA (Figure 4).

**Fig. 4.**
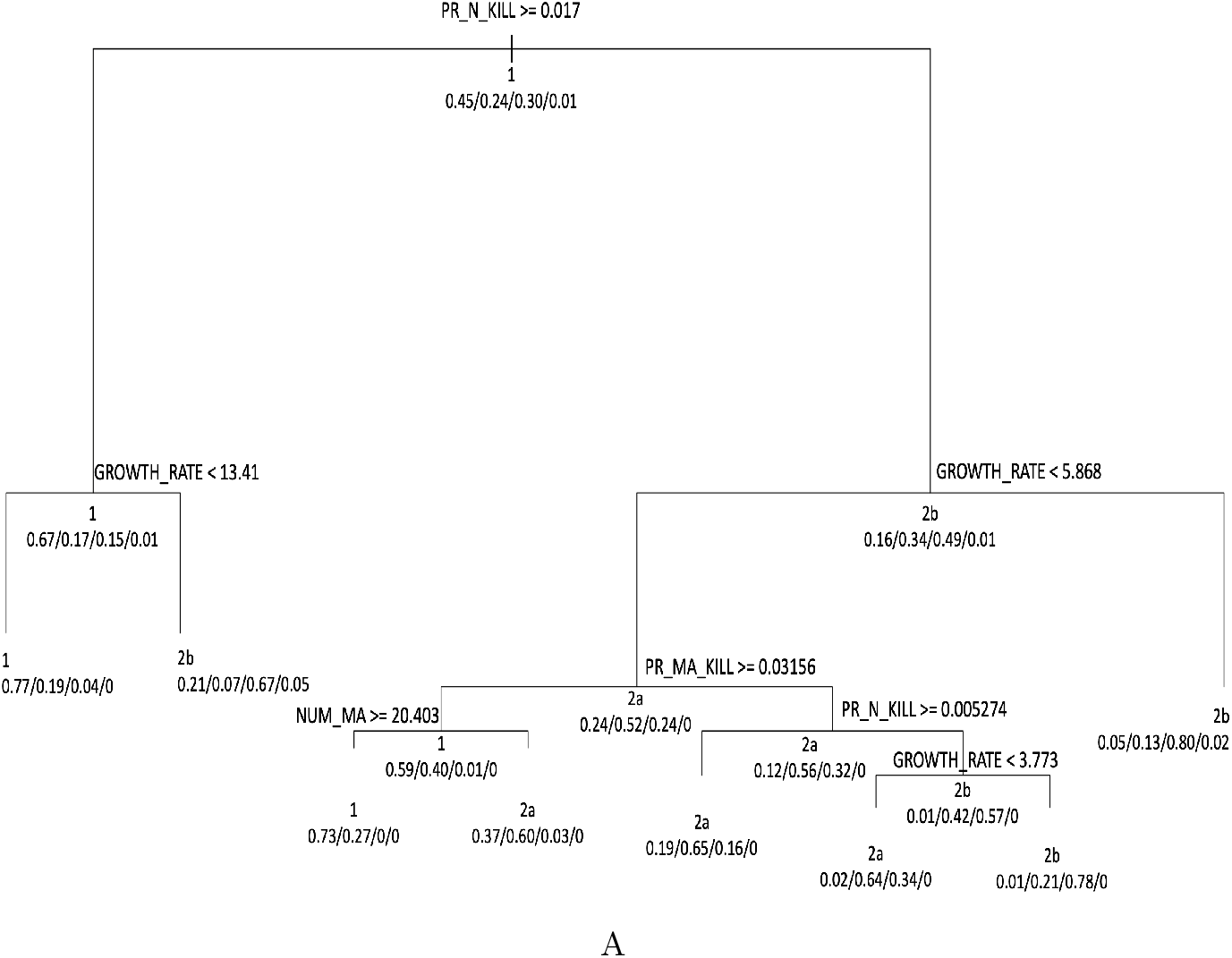
Classification tree. The classification tree found the parameter values that best parse the virtual patients into the four clusters defined in Figure 3. The percentage value range is 0-1. The sets of four numbers beneath each branch are the relative frequency of each cluster in that branch.

One can see in Figure 4 that the dominant parameters are the neutrophil killing rate, intrinsic growth rate, and to a lesser degree, the monocyte killing rate. Classification trees can indicate which parameters are the most important by measuring how much each parameter contributes to node purity. We confirmed that these three parameters (neutrophil killing rate, intrinsic growth rate, and monocyte killing rate) are indeed the dominant ones. Similarly, in Scenario 2, granule half-life, *k_d_*, and to a lesser degree, intrinsic growth rate are the dominant parameters (not shown, Figures A3).

Since we have only three dominant parameters, we can plot the virtual patients in a 3D space. That will allow us to see how the four clusters are distributed. In Figure 5A, each point represents a virtual patient. The x-axis is the log of that patient’s intrinsic growth rate, the y-axis is the log of that patient’s monocyte killing probability, and the z-axis is the log of that patient’s neutrophil killing probability. We colored each patient according to their cluster (1, 2A, 2B, or 2C). We used log base ten because different patients have parameter value differences of up to two orders of magnitude.

**Fig. 5.**
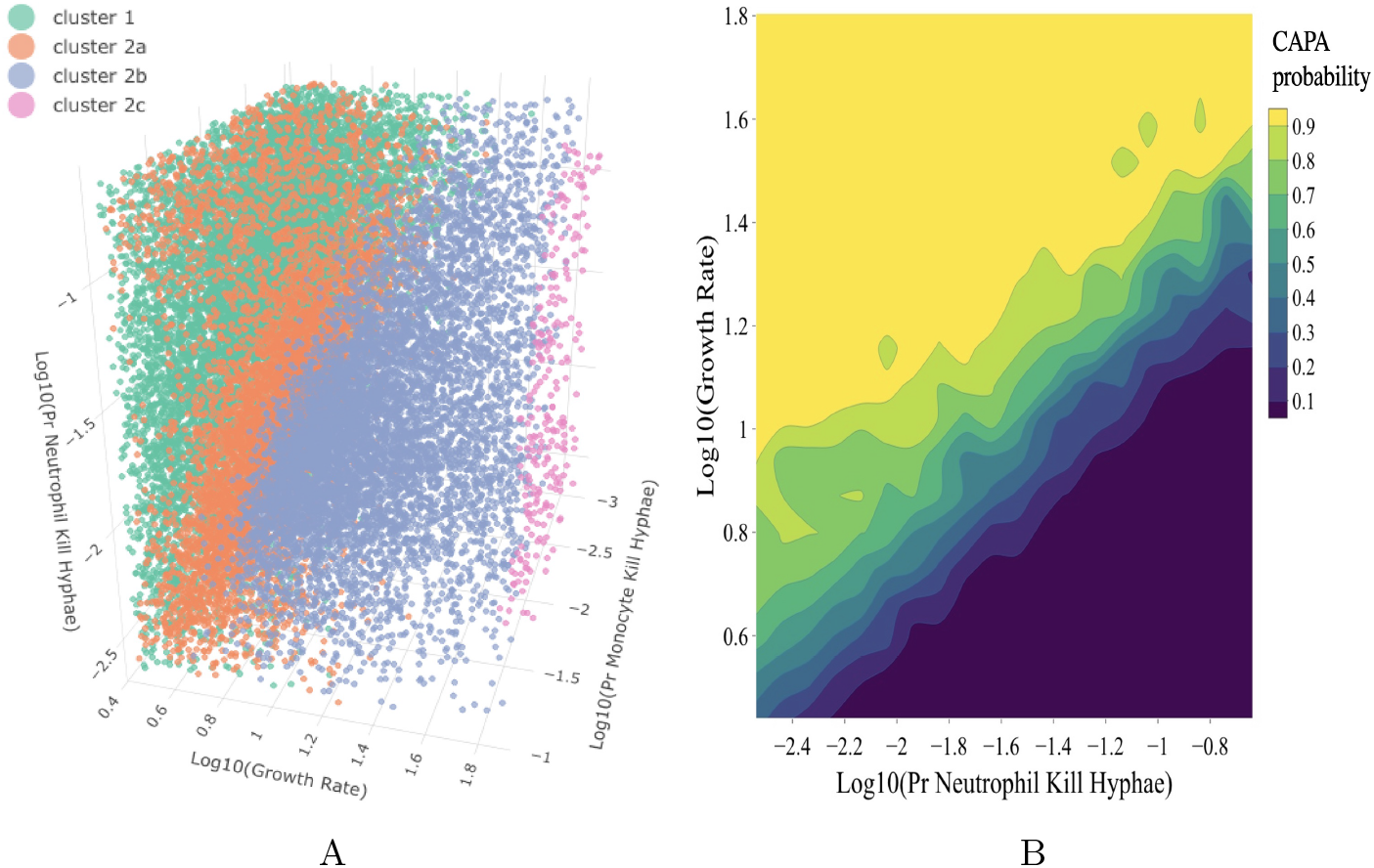
Distribution of clusters of patients and CAPA probability in the 3D and 2D spaces formed by the most important parameters according to the classification tree (Figure 4). Figure 5A virtual patients plotted in 3D space and colored according to their cluster. The x-axis is the log of patients’ intrinsic growth rate, the y-axis is the log of their monocyte killing probability, and the z-axis is the log of the neutrophil killing probability. Figure 5B shows the CAPA probability as the intrinsic growth rate and neutrophil killing probability.

As shown in Figure 5A, Clusters 1 and 2A are interspersed. This is to be expected because both these clusters represent patients for whom the infection cleared (Cluster 2A is the negative part of peak 2). Cluster 2C is segregated in the high intrinsic growth rate region. However, it is noteworthy that some patients in Cluster 2B are very similar to patients in Cluster 2A, in that they have similar sets of parameters that characterize their infection.

To better explore this transition between patients that clear the infection (Clusters 1 and 2A) and those that do not (Clusters 2B and 2C), we computed how the probability of developing CAPA changes as parameters vary (Figure 5B). We restrict ourselves to plotting the CAPA probability across the two most influential parameters. We plot how a virtual patient’s probability of developing CAPA changes as the intrinsic growth rate and the neutrophil killing rate change (Figure 5B). As shown in Figure 5B, the dividing line along which a patient may either clear the infection or develop CAPA cuts the graph across the diagonal. Both parameters (intrinsic growth rate and neutrophil killing probability) are essential in determining the patient’s outcome: a high intrinsic growth rate can be compensated for, in part, by a high killing rate and vice versa. In Scenario 2, we found a similar transition between patients that develop CAPA and those that do not when we vary the parameters of intrinsic growth rate and granule *k_d_* (Figure A3B). This shows that our conclusions are similar in both scenarios.

An essential question in Figure 5, especially 5B, is whether using only two parameters is meaningful. To answer that question, we divided the virtual patients into two subsets. The first subset of 20,000 was used to train classification trees, while the second subset of 4,000 was used to test its predictive power. We also used ten-fold cross-validation and measured the F1 score, as in Table 2. The F1 score combines the precision and recall into a single metric, and it is primarily used to compare the performance of a classifier. We found that the reduction from all five parameters to the top three (GROW_RATE, PR_N_KILL, and PR_MA_KILL) does not affect the predictive power of the classification trees, while the reduction to the top two (GROW_RATE and PR_N_KILL) has only a minor effect.

**Table 2.**
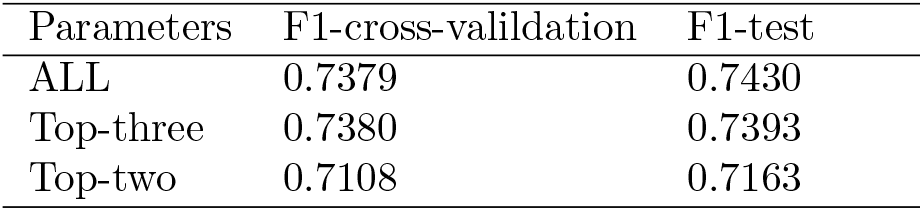
The predictive power of parameters. The predictive power of classification trees was evaluated in cross-validation and test sets when these trees were trained using all five parameters, the top three parameters (GROW_RATE, PR_N_KILL, and PR_MA_KILL), or the top two (GROW_RATE and PR_N_KILL). Column 1 parameters used in the model; Column two F1 score in cross-validation; Column 3 F1 score in the test set.

| Parameters | F1-cross-validation | F1-test |
| --- | --- | --- |
| ALL | 0.7379 | 0.7430 |
| Top-three | 0.7380 | 0.7393 |
| Top-two | 0.7108 | 0.7163 |

Results from previous figures have shown that the observed growth rate is highly dependent on the intrinsic growth rate. We divided the virtual population into ten bins of similar intrinsic growth rates and computed the square of the partial rank correlation of the parameters and observed growth rate (Figure 6). Figure 6A shows the variation of the square of the correlation between neutrophil killing probability and observed growth rate. At the same time, 6B shows the variation of the square of the correlation between monocyte killing probability and observed growth rate. The partial correlation between the other parameters and the observed growth rate was not substantial (results not shown).

**Fig. 6.**
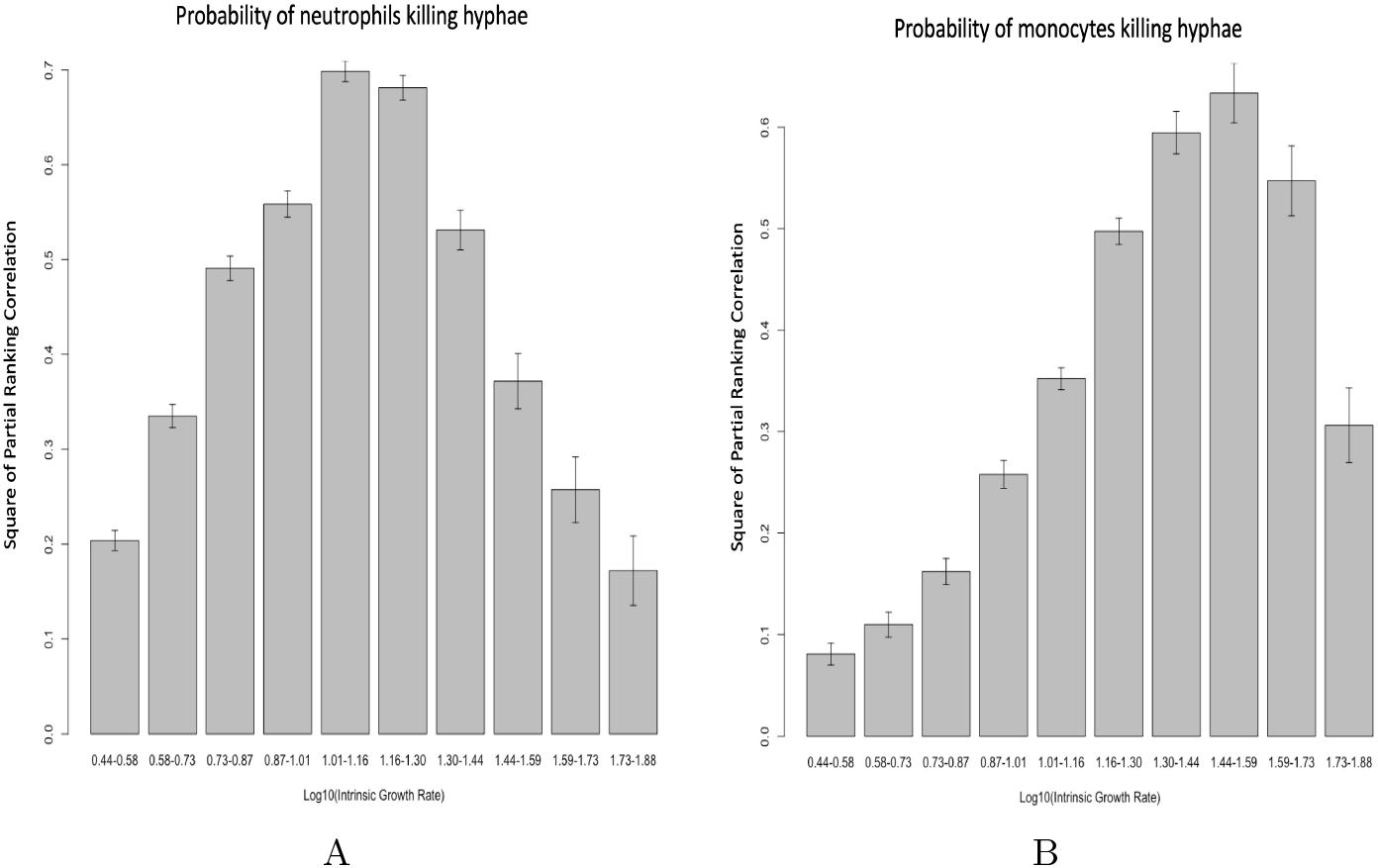
Variation of the square of the correlation *r*^2^ between the critical model parameters and the observed growth rate, with the variation of the intrinsic growth rate. We started with a dataset of 24,000 virtual patients generated with LHS and then partitioned this dataset by intrinsic growth rate. The x-axis shows the range of the intrinsic growth rate in each partition. We then calculate the correlation between parameters and the observed growth rate in each partition. Figure 6A: Variation of *r*^2^ between the neutrophil killing rate and observed growth rate. Figure 6B: Variation of *r*^2^ between monocyte killing rate and observed growth rate. Error bars are standard deviations calculated with bootstrap.

## 4 Discussion

*Aspergillus fumigatus* is an opportunistic mold that infects immunocompromised hosts, chiefly those with impaired neutrophil response, but also patients with normal neutrophils but impaired cell-mediated immunity [18]. However, Lay, C, and Yu, W 2021 [3] found that in patients with CAPA the conventional risk factors for invasive aspergillosis are not present. They found that CAPA patients were mostly older men suffering from diabetes, obesity, hypertension, or cardiovascular disease. These conditions may help explain the onset of CAPA, but are likely not sufficient by themselves. We hypothesized that COVID-19 infection may exhaust the immune system, and at the same time the destruction of lung epithelium overloads the fungus with nutrients. Therefore we propose a model where the coinfection negatively affects the default parameters we inherited from [7]. We tested this hypothesis using a simulated virtual patient population generated by our computational model.

Before we can draw conclusions from the model, it is critical to discuss the parameters. Like our previous one, this model was parameterized with literature data. Therefore it does not fit any particular dataset exactly. However, it reproduces biological data within the limit of biological variance itself. That was the case with the several datasets with which we compared our model in our previous paper [7]. We note that it is possible, to fit this model to particular data sets.

Another point that is worth discussing is identifiability. Because of the way parameters were obtained from the literature (see [7]), identifiability is not an issue here. As an example, take cytokines *k_d_*, secretion rate, and half-life. If one tries to fit these parameters from a time series of cytokines in BAL or lung homogenate in an infection, likely, they will not be identifiable. However, we got these parameters from experiments designed to measure each one individually.

Concerning the experiments we conducted, our simulations show that 31% of the virtual patients in Scenario 1 and 17% in Scenario 2, who were exposed to *A. fumigatus*, developed CAPA. That is qualitatively consistent with data from [4] that show 10% of patients in the ICU developed CAPA and with data from [19] that show that 15% of patients in the ICU developed CAPA (Figure A1). Moreover, Figures A4 and A5 suggest that the kinds of parameter variations proposed in this model (i.e., change in *A. fumigatus* virulence and neutrophil activity) exist in the general population. On the other hand, the discrepancy, chiefly in Scenario 1, can be explained by the fact that most patients hospitalized with COVID-19 infection are presumably not exposed to *A. fumigatus*.

Results from Figure A1 show that Scenario 2 seems to agree more with epidemiological data, but it also shows a discrepancy between the two scenarios. This highlights the fundamentally different dynamics of neutrophils killing by direct contact or secreting granules that kill hyphae in the neighborhood. Both scenarios were made to fit the simulations in [7]. However, the two scenarios diverged when we varied the conditions, albeit still behaving in similar ways.

Our model suggests that both reducing neutrophil activity and increasing fungal intrinsic growth rate might be necessary for CAPA to emerge (Figures 4 and 5). Tappe, B *et al*. 2022 [20] found that neutrophils of COVID-19 patients had reduced ROS production qualitatively similar to the reduction in neutrophils’ killing ability in simulated CAPA patients in our model (Figure A4). At the same time, Grunwell, JR *et al*. 2018 [21] found that neutrophils treated with airway supernatant of patients with acute respiratory failure due to lower respiratory tract viral infection had altered surface markers. Moreover, when the neutrophils were treated with airway supernatant in patients with bacterial and viral coinfection, they had decreased respiratory burst and bactericidal response. The mechanism is not well understood. However, Verweij, PE *et al*. 2020 [22] reported that influenza might suppress neutrophil oxidative burst, causing temporary disease status resembling chronic granulomatous disease. Arastehfar, A *et al*. 2020 [23] indicate that collateral effects of antiviral immunity may, paradoxically, contribute to an inflammatory environment that favors secondary infections such as CAPA. On the other hand, the drugs used to treat serious COVID-19 infection, such as Tocilizumab and dexamethasone, can also hinder the immune response against *A. fumigatus* [24]. These findings support the hypothesis that the immune response against *A. fumigatus* may be hampered in patients with serious COVID-19 infection.

Concomitantly, Escobar, N *et al*. 2016 [25] showed that pneumocytes cocultured with *A. fumigatus* inhibited fungal germination and growth. In contrast, Rodrigues, AG *et al*. 2005 [26] found that exposing *A. fumigatus* to albumin increases its germination and growth rate. Hsu, JL *et al*. 2018 [27] gives a persuasive argument for hemorrhaging increasing the rate of *A. fumigatus* invasion. Likewise, Michels, K *et al*. 2022 [28] found that heme increases fungal growth (Figure A5). While Figure A5B shows only the effect of heme *in-vivo* Michels’s paper [28] gives compelling evidence that heme iron and perhaps heme ring favors *A. fumigatus* growth *in-vitro* and *in-vivo*.

In our previous model, iron concentration was a bottleneck reducing the observed growth rate [7]. Hemorrhage caused by previous COVID-19 infection may help to overcome this bottleneck, according to the evidence provided by [27, 28]. Furthermore, the iron concentration (non-heme) may be higher in serum than in the alveolar space [29]. Arastehfar, A *et al*. 2020 [23] argue that the leading risk factor for CAPA includes severe lung damage during the course of COVID-19 and the presence of comorbidities such as structural lung defects. These findings support our hypothesis that lung epithelium destruction and hemorrhage favor *A. fumigatus* growth: once the pneumocytes are killed, *A. fumigatus* is exposed to a nutrient-rich environment and can grow faster.

Figure 5 suggests that the onset of CAPA is regulated chiefly by the intrinsic fungal growth and neutrophil killing rates. The CAPA probability increases as the intrinsic growth rate increases or as the neutrophil killing probability decreases. In the alternative neutrophil action scenario (Figure A3), we can make a similar observation with the granule *K_d_* and intrinsic growth rate (Figure A3B). A moderate increase in the intrinsic growth rate and decrease in neutrophil activity can also lead to CAPA. Alternatively, a high increase in intrinsic growth rate or a steep decrease in neutrophil activity could also lead to the same outcome. Both scenarios we tested (i.e., neutrophils killing by direct contact or by secreting granules) support this conclusion. We found the first hypothesis to be more plausible (i.e., moderate change). However, Grunwell, JR *et al*. 2018 [21] found a substantial reduction in neutrophil killing ability that could support the second hypothesis (i.e., high change in neutrophil killing probability). Whether their result generalizes to fungal coinfection remains open.

Figure 6 shows that neutrophils are more efficient in controlling the infection in the case of a moderate intrinsic growth rate. This corroborates the previous results. It may also reflect how identifiable these two parameters are. Figure 6 can be divided into two: the center and the tails. In the tails, the leukocytes always win or lose the battle independent of the value of the intrinsic growth rate. But that means that the observed growth rate in these regions is not strongly correlated with leukocyte activity. The center is where there is an exchange: as the intrinsic growth rate increases, the neutrophil and monocyte activity has to increase to keep the fungus in check. That means that these two parameters are not identifiable from the point of view of the observed growth rate. That reinforces the previous conclusion that CAPA may be caused by decreased leukocyte activity, increased *Aspergillus fumigatus* growth, or both.

Figures 5B and A3B, however, suggests that CAPA might be caused by both instead of only one of the causes. The tails and the center in Figure 6 roughly relate to the plateaus and the dividing line in Figure 5B and A3B. Note that there are only narrow areas of intrinsic growth rate that are entirely independent of neutrophil activity. That is, the dividing line in graphs 5B and A3B cuts the plot along the diagonal.

Interestingly, the work of Delliere, S *et al*. 2021 [30] analyzed several clinical factors in COVID-19 patients with and without CAPA. Treatment of COVID-19 patients with the antibiotic azithromycin was associated with an increased risk of CAPA. In prior literature, azithromycin has been shown to attenuate neutrophil oxidative burst[31]. In our model, this would translate into a decreased neutrophil-killing probability. This fits with our finding that this parameter is paramount for CAPA establishment.

On the other hand, Delliere, S *et al*. 2021 [30] found similar percentages of neutrophils and macrophages in CAPA and non-CAPA patients. Likewise, Xu, J *et al*. 2021 [32] found that patients with fewer than 1.5 ×10^9^ neutrophils/L had no increased risk of developing CAPA. That resonates with our finding that the number of neutrophils and monocytes had a low correlation with the observed growth rate (Table A4) and was a poor predictor of CAPA (Figure trees).

We conclude that an increase in the favorability of the lung environment, likely due to an increased supply of nutrients and decreased effectivity of neutrophils to kill hyphae, is the most likely explanation for CAPA. To make our findings more robust, we tested two possible scenarios of how neutrophils kill hyphae (Supplementary Material). We found that the conclusions are the same, independent of the scenario.

## Ethics approval

The manuscript complies with the ethical standards of the Journal for Mathematical Biology.

## Consent to participate

Not applicable

## Consent for publication

All authors have approved the manuscript

## Availability of data and materials

Simulation data available on request

## Code availability

Code available upon request

## Authors’ contributions

The study was conceived and carried out by HALR, with scientific input from the other authors. RL an BM provided funding for the project and contributed to the writing of the manuscript.

## Appendix A Supplementary Material

**Table A1.**
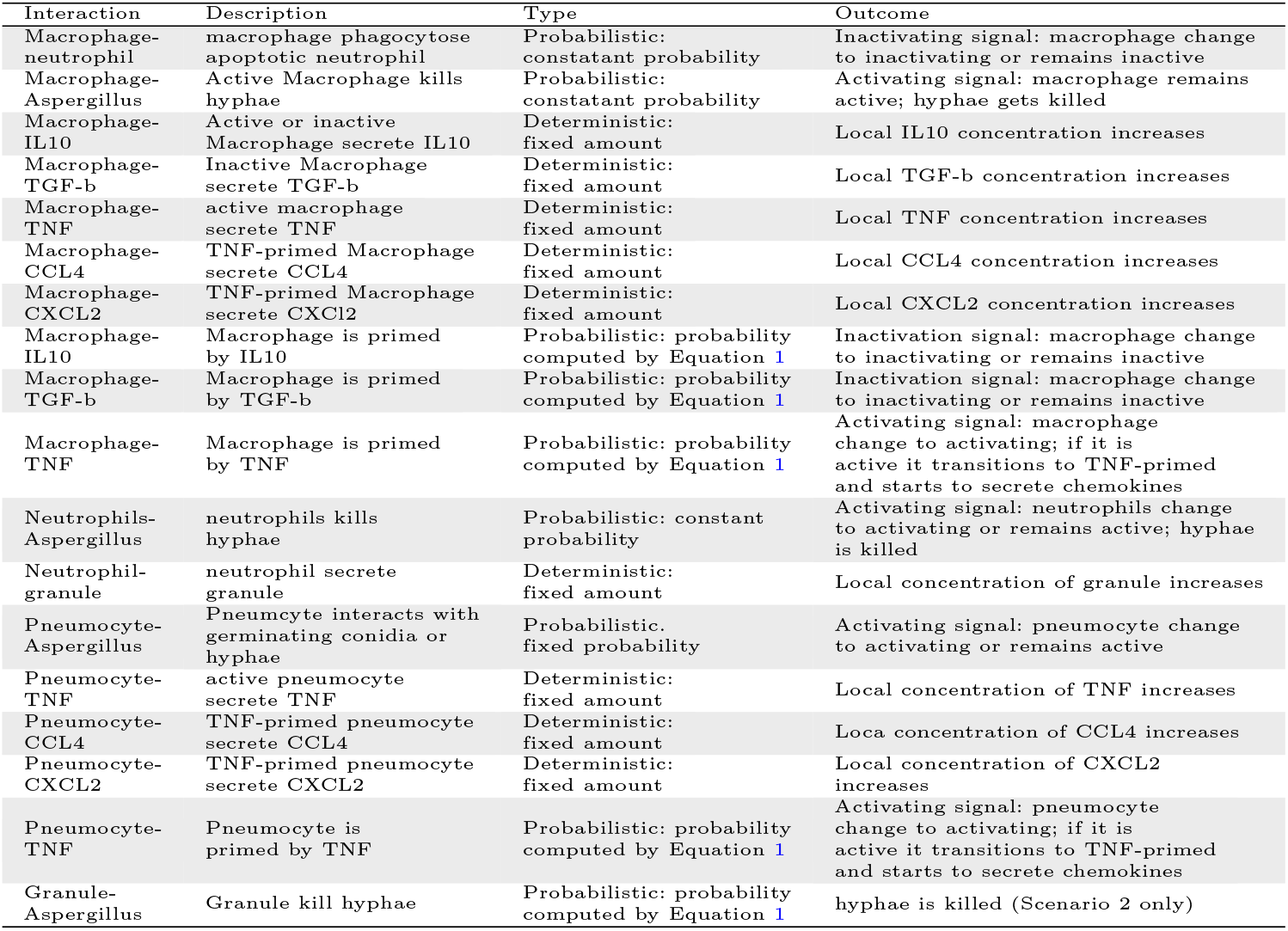
Table depicting the rules of how agents and field variables interact. The first column shows the two entities involved in the interaction. The second column briefly describes the interaction; the third indicates whether it is deterministic or probabilistic and how the probability is computed. The fourth column describes the outcome of the interaction in case of success. Note that two entities (Column 1) can interact in more than one way.

### A.1 Rules

Table A.1 shows the rules used in the model, all taken from [7]. That is, the rules in this model are a subset of the rules in our previous model. The reduction in the number of rules is related to the fact that we are not simulating iron explicitly. Second, we start the model with already germinated conidia (i.e., small hyphae). Therefore, rules such as the interaction of leukocytes with conidia and conidia germination are unnecessary. Table A.2 shows the parameters in the model; again, all taken from [7], except for the number of leukocytes. Subsequently, we provide a brief rationale for all the parameters with references.

### A.2 Parameters

The *k_d_* of TNF, IL10, TGF, CCL4, and CXCL2. The *k_d_* values from these molecule receptors are reported by [33–50]. In cases where we have more than one value we used the median.

**Table A2.** Parameters used in the model. The first column gives the parameter name; the second a short description; the third the parameter value; and the forth a bibliographic reference. Note that apart from the number of leukocytes, all parameters were inherited from our previous model.

| Parameter | Description | Value | Reference |
| --- | --- | --- | --- |
| TNF_QTTY | TNF secretion quantity | $1.46 \times 10^{-20} \text{ mol} \times \text{cell}^{-1} \times \text{h}^{-1}$ | [7] |
| IL10_QTTY | IL-10 secretion quantity | $6.97 \times 10^{-22} \text{ mol} \times \text{cell}^{-1} \times \text{h}^{-1}$ | [7] |
| TGF_QTTY | TGF-β secretion quantity | $1.01 \times 10^{-21} \text{ mol} \times \text{cell}^{-1} \times \text{h}^{-1}$ | [7] |
| CCL4_QTTY | CCL4 secretion quantity | $1.79 \times 10^{-20} \text{ mol} \times \text{cell}^{-1} \times \text{h}^{-1}$ | [7] |
| CXCL2_QTTY | CXCL2 secretion quantity | $1.11 \times 10^{-19} \text{ mol} \times \text{cell}^{-1} \times \text{h}^{-1}$ | [7] |
| $k_d$_TNF | $k_d$ TNF receptor | 326 pM | [7] |
| $k_d$_IL10 | $k_d$ IL10 receptor | 140 pM | [7] |
| $k_d$_TGF | $k_d$ TGF-β receptor | 26.5 pM | [7] |
| $k_d$_CCL4 | $k_d$ CCL4 receptor | 180 pM | [7] |
| $k_d$_CXCL2 | $k_d$ CXCL2 receptor | 91.667 pM | [7] |
| D | diffusion coefficient | $850 \mu\text{m}^2/\text{min}$ | [7] |
| r | elongation rate | $40 \mu\text{m}/\text{h}$ | [7] |
| PR_BRANCH | branch probability | 25% | [7] |
| MV_RATE | leukocyte move rate | $1.44 \mu\text{m}/\text{min}$ | [7] |
| N_H_KILL | probability of neutrophil killing hyphae | 22.71% | [7] |
| MA_H_KILL | probability of macrophage killing hyphae | 9.81% | [7] |
| E_INT | epithelial cells-Aspergillus interaction rate | 4.49% | [7] |
| L | molecule half-life | 1h | [7] |
| TURNOVER_RATE | molecule exchange rate between lung and whole body serum | $0.1823 \text{ h}^{-1}$ | [7] |
| MA_HALF_LIFE | macrophage half-life | 24h | [7] |
| N_HALF_LIFE | neutrophils-half-life | 6h | [7] |
| SEPTAE_L | septa length | $40 \mu\text{m}$ | [7] |
| ITER_CH_STATE | iterations for cell to change state | 60 | [7] |
| ITER_TO_REST | iterations for cell to return to resting state | 180 | [7] |
| NUM_MA | number of macrophages | 960 | [12] |
| NUM_N | number of neutrophils | 360 | [12] |
| NUM_P | number of pneumocytes | 640 | [7] |

The TNF, IL10, TGF, CCL4, CXCL2 secreting rates come from [51–80]. For more detail see [7].

Diffusion rate of cytokines is reported by [81, 82].

Half-life of cytokines is reported by [83–89].

This movement rate can be obtained from Khandoga, AG et al. 2009 [90]. The value, 1.44 *μm/min*, is conservative compared to other sources. Pollmacher J, & Figge MT, 2014 [91] uses a movement rate of 2-6 4 *μm/min*, for instance. Nevertheless, the rate used here must be considered a phenomenological movement rate. In the real lung, leukocytes may not move in a straight line but along the alveolar curved surface. That is the case in the Pollmacher J, & Figge MT, 2014 [91] model.

Growth rates come from papers that report hyphal length over time [12, 25, 92, 93], while branching probability was based on the hyphal growth unit length. This gives an estimate of how many branches per septum there are [92, 94].

Monocyte and neutrophil killing probabilities are extrapolated from the killing rate of *in vitro* experiments from [10, 11, 95–98]. Likewise for the pneumocyte interaction rate [89]. For more details see [7].

Turnover rate comes from the difference between lung and serum IL-6 concentration [99]. For more details see [7].

Monocyte and neutrophil half-life is reported in [100, 101].

Septae length is reported in [102, 103].

The time that cells need to change status (T CHANGE and T REST - Figure 2) were based on *in vitro* reports [104].

Number of macrophages and monocytes comes from mice infected with influenza [12]. While the number of pneumocytes comes from [105].

**Table A3.** Model parameter ranges for the alternative scenario. The table shows the ranges in which we sample parameters (maximum and minimum value). Column 1: parameter name; Column 2: description; Column 3: parameter range; Column 4: parameter reference value; Column 5: rationale for the values in Column 4. The values taken from [7] are the default values in our previous work. The values for GROWTHRATE, PR_MA_KILL, NUMMA, and NUM_N are the same as in Table 1

| Parameter | Description | Parameter range | Reference Value | Ref |
| --- | --- | --- | --- | --- |
| GROWTH_RATE | intrinsic growth rate | 2.75-80 $\mu\text{m}/\text{h}$ | 40 $\mu\text{m}/\text{h}$ | [7] |
| PR_MA_KILL | probability of monocytes to kill hyphae | 0.099-9.9% | 9.9% | [7] |
| $K_d$-GRANULE | $K_d$ of the reaction between granule and hyphae | $1 \times 10^8 - 1 \times 10^{10}$ | $1 \times 10^{10}$ | [7] |
| GRANULE_HALF_LIFE | granule half-life | 0.033 - 3.33 h | 0.67h | [7] |
| NUM_MA | number of monocytes | 360-960 cells | 960 cells | [16] |
| NUM_N | number of neutrophils | 360-960 cells | 360 | [16] |

#### A.3 Supplementary Results

This supplementary material contains results from Scenario 2, sensitivity analysis, and comparisons of simulated results with the literature.

**Table A4.** Table with the sensitivity analysis (SA) of the parameters used to create the virtual population. The parameter range is shown in Table A3. First column parameters description; Column 2 partial rank correlation for Scenario 1; Column 3 partial rank correlation for Scenario 2.

| Parameter | Scenario-1 | Scenario-2 |
| --- | --- | --- |
| GROWTH_RATE | $0.5538 \pm 0.0050$ | $-0.0317 \pm 0.0061$ |
| PR_N_KILL | $-0.5497 \pm 0.0042$ | NA |
| PR_MA_KILL | $-0.3170 \pm 0.0061$ | $-0.0788 \pm 0.0052$ |
| NUM_N | $-0.1114 \pm 0.0062$ | $-0.1093 \pm 0.0050$ |
| NUM_MA | $-0.0780 \pm 0.0064$ | $-0.0535 \pm 0.0050$ |
| GRANULE_HALF_LIFE | NA | $0.5929 \pm 0.0038$ |
| $K_d$_GRANULE | NA | $-0.3809 \pm 0.0049$ |

**Fig. A1.**
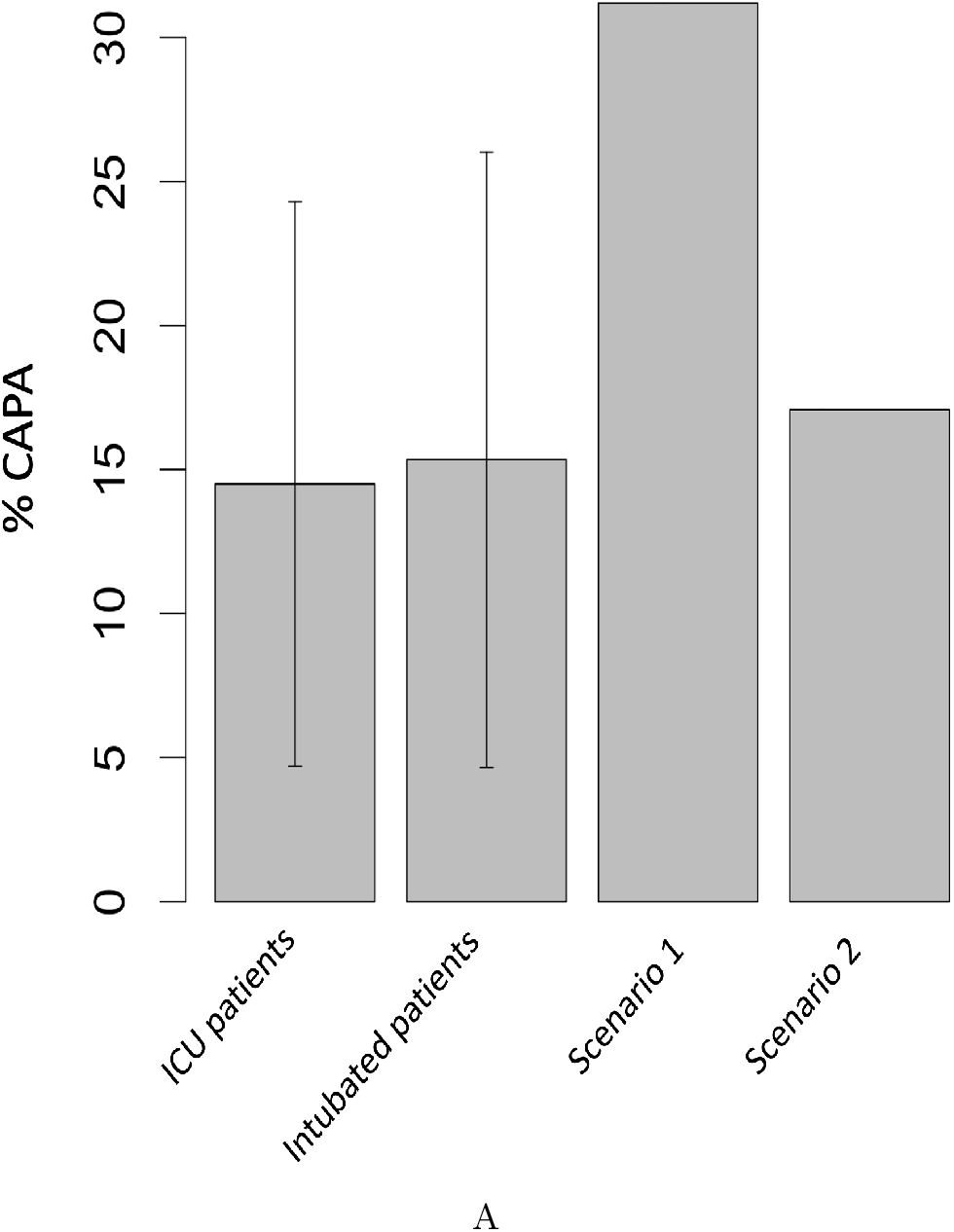
Comparison between epidemiological data on CAPA [19] and virtual epidemiology. ICU patients and intubated patients refer to epidemiological data from [19]. Scenarios 1 and 2 are simulation data. Error bars represent standard deviation.

**Fig. A2.**
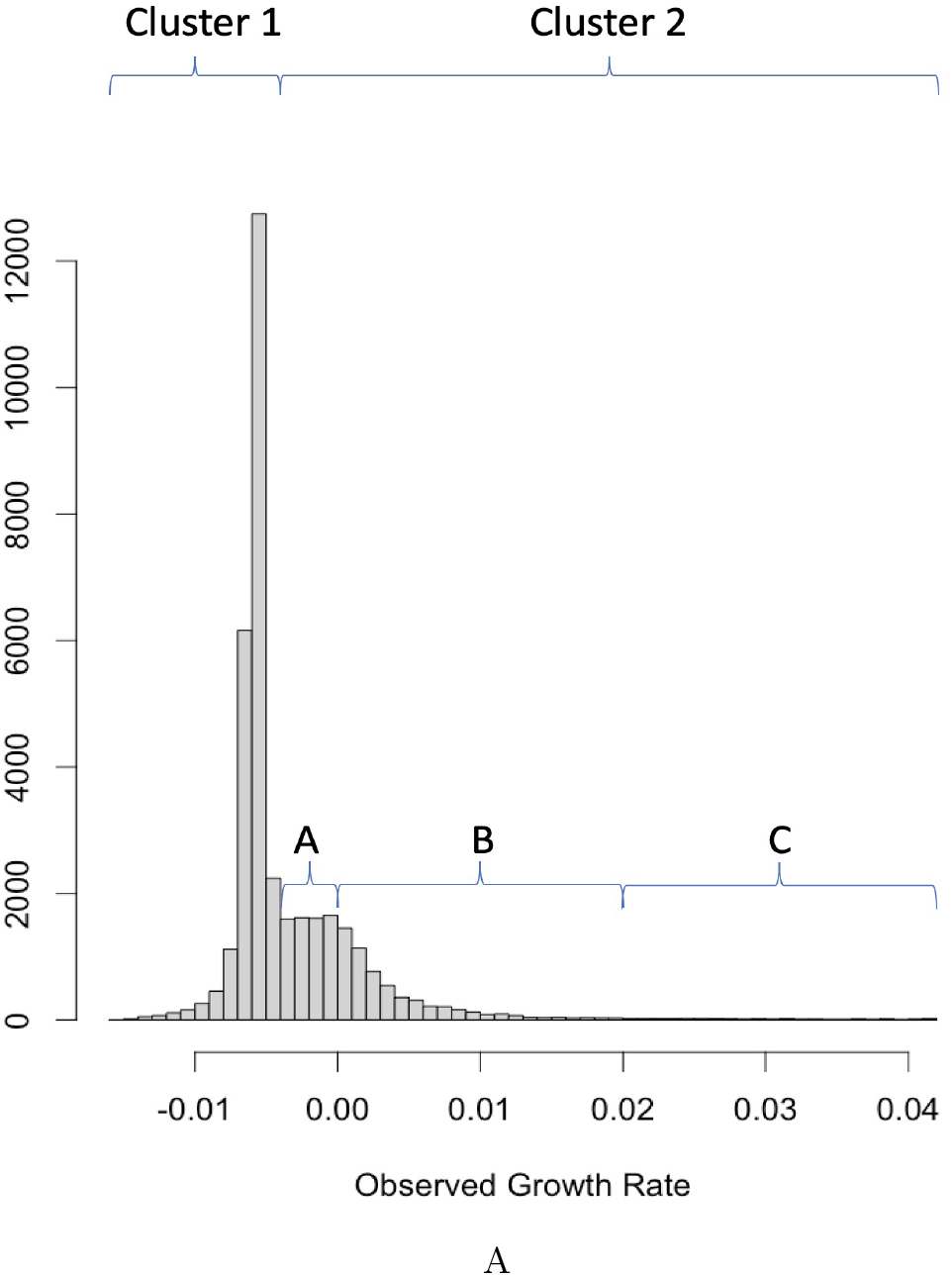
Distribution of observed growth rate in the alternative scenario where neutrophils do not need direct contact to kill hyphae. We divided virtual patients into two clusters: the prominent peak on the negative side (Cluster 1) and the secondary peak (Cluster 2). We then subdivided Cluster 2 into three subclusters: 2A is the negative part, 2B is the positive part excluding outliers, and 2C is the long tail of outliers.

**Fig. A3.**
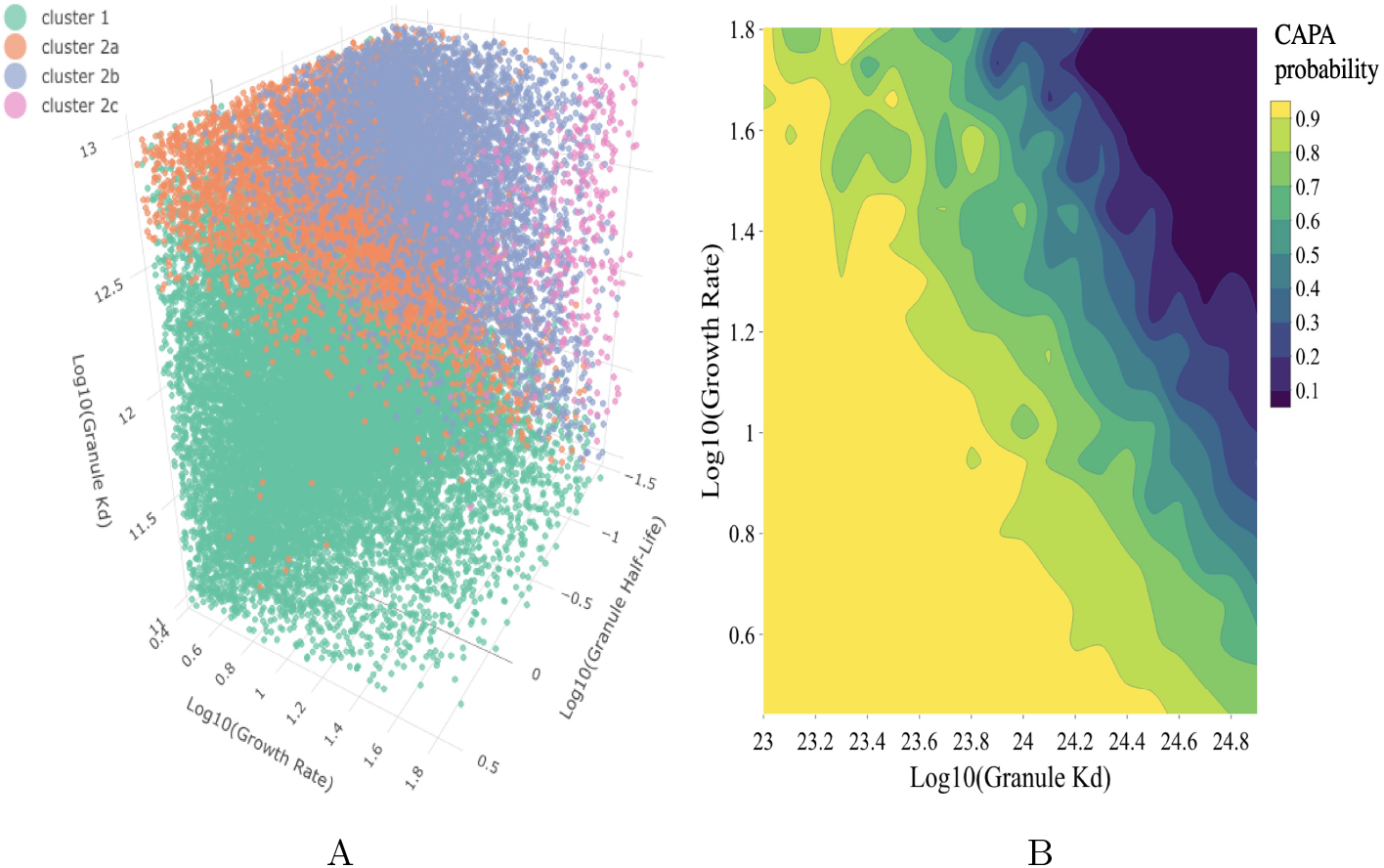
Plots in 3D space of the most important parameters according to the classification tree (not shown) in the alternative scenario where neutrophils do not need direct contact to kill hyphae. Figure A3A Virtual patients plotted in 3D space. The x-axis is the log of patients’ intrinsic growth rate, the y-axis is the log of their neutrophil granule *k_d_*, and the z-axis is the log of the neutrophil granule half-life. Figure A3B shows the CAPA probability as the intrinsic growth rate and neutrophil granule *k_d_* change.

**Fig. A4.**
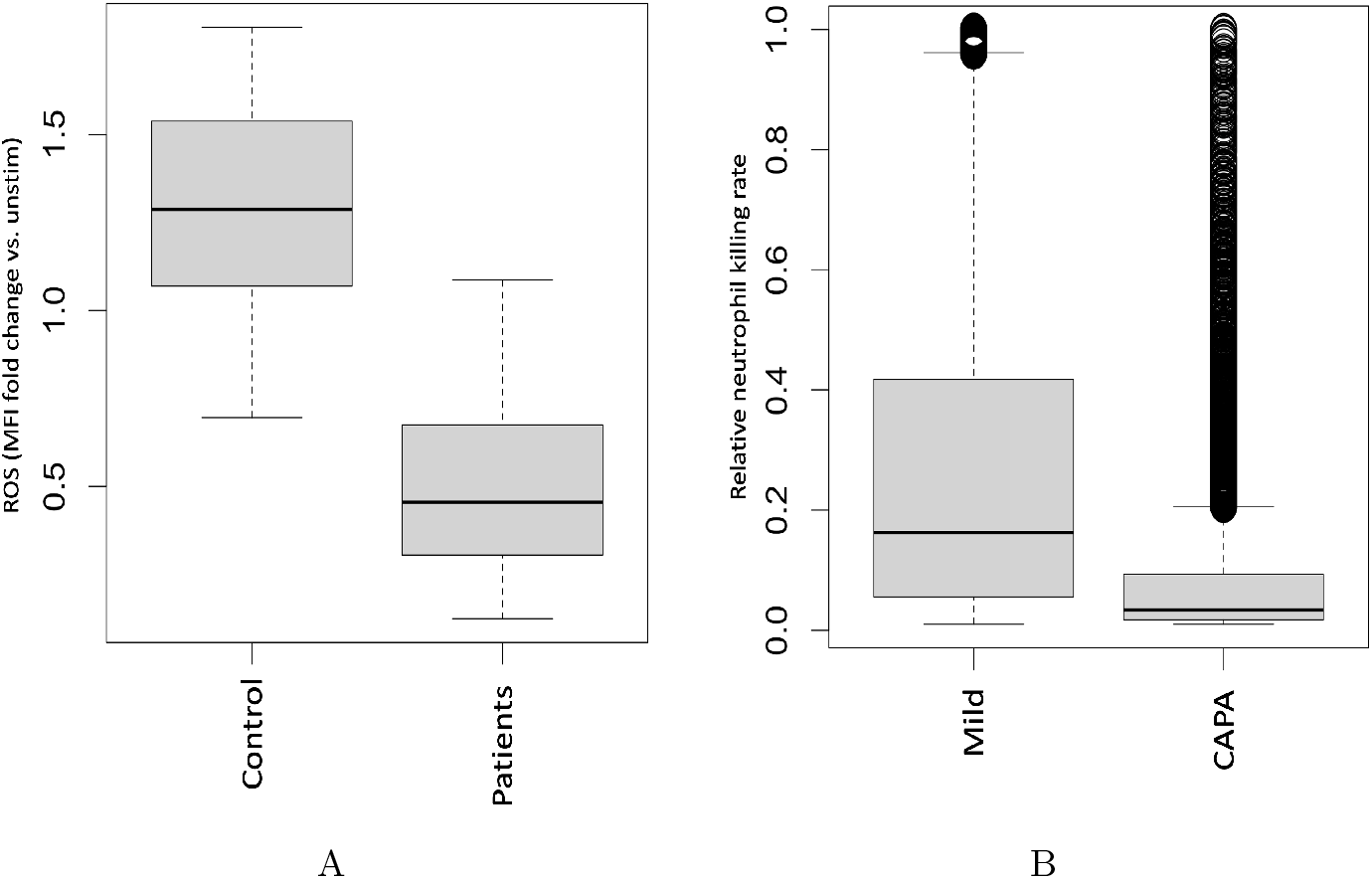
Immune system inhibition by SARS-CoV-2. A: decrease in ROS production in patients with SARS-CoV-2 [20]. B: decrease in neutrophil killing ability in virtual patients with CAPA relative to patients without CAPA (Scenario 1).

**Fig. A5.**
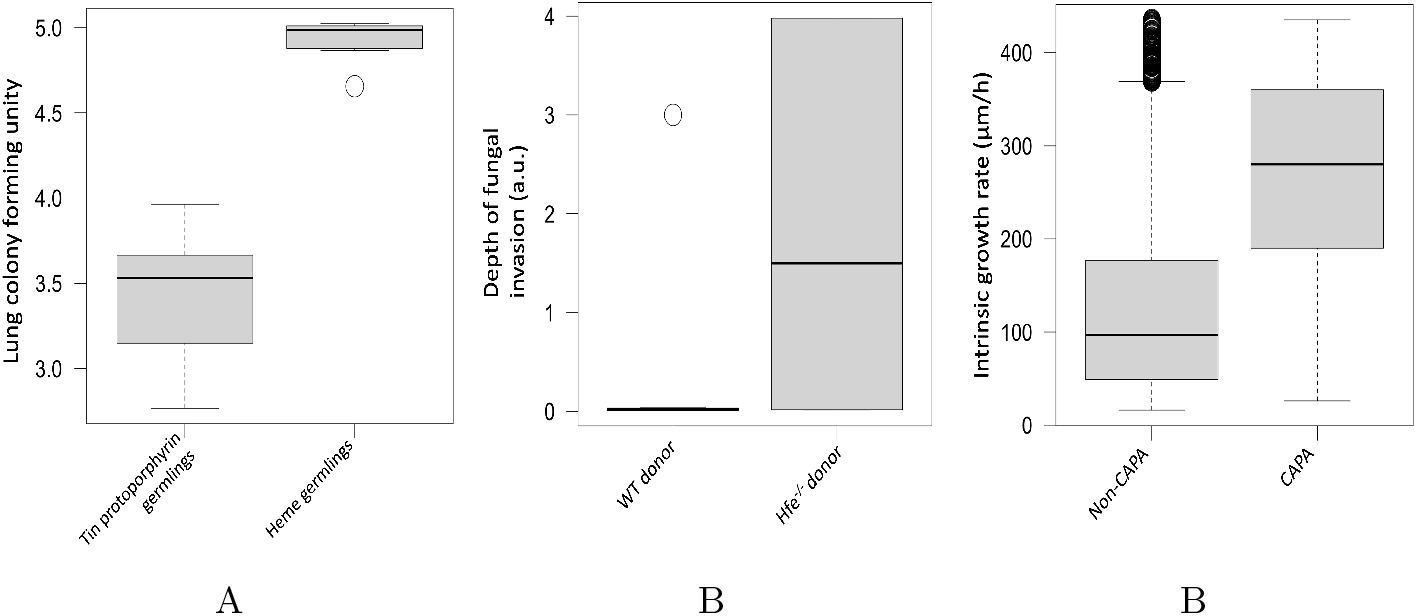
Hemorrhage leads to increased *A. fumigatus* growth *in-vivo* due to iron availability. A: Mice treated with heme and infected with A. *fumigatus* had more lung colony-forming units than litter mates treated with Tin protoporphyrin. B: Lung transplant causes micro-hemorrhage that predisposes mice to fungal invasion due to iron availability. Hemochromatosis mice Hfe-/-mice had more fungal invasion control when infected with *A. fumigatus* after receiving a lung transplant. C: simulated CAPA virtual patients (Scenario 1) had higher intrinsic growth rates, similar to what A and B suggest for mice with hemorrhage.

